# Assessing natural selection during range expansions: Insights from a spatially explicit ABC study

**DOI:** 10.1101/391110

**Authors:** Ricardo Kanitz, Samuel Neuenschwander, Jérôme Goudet

## Abstract

For at least 40 years now, evolutionary biologists have discussed the relative roles of natural selection and genetic drift in shaping the genetic composition of populations. Range expansions are of particular interest in this discussion: They normally occur over environmental gradients allowing local adaptation to take place, but the demographic properties of these expansions also potentiate genetic-drift effects, which may in turn randomly generate extreme changes in allele frequencies as populations expand in territory and numbers (i.e. allele surfing). Here, we address the detection and measurement of selection in such scenario using simulations. We mimic a range expansion over a variable selective gradient where individuals have in their genomes both loci that are neutral and loci determining a quantitative trait subject to selection. The responsiveness of summary statistics to the selective pressure is then assessed, and estimates of the selective pressure are made – based on these statistics – with approximate Bayesian computation (ABC). We observe that statistics related to isolation-by-distance patterns present a strong response to selection. This response can be used in ABC to estimate the strength of selection acting on the simulated populations with very reliable measures of estimability, regardless of the genetic architecture underlying the selected trait. Furthermore, these estimates are robust to noise produced by other genetic and demographic parameters such as heritability, mutation, migration and population-growth rates. This approach of taking into account the spatial dimension of differentiation in quantitative traits offers a promising avenue of investigation about the role of natural selection in range-expansion scenarios, with possible implementations in the study of natural cases, as well.

## Introduction

The opposition between selectionism and neutralism is one of the most significant debates in evolutionary biology (Ewens 1977; Kimura 1984; Hey 1999; Nei 2005). Ultimately, the question relies on which kind of processes (neutral or selective) lead to the majority of patterns observed in nature. Even though reconciliatory ideas have been proposed (Wagner 2008), the dilemma regarding selection vs. neutrality still endures in different contexts of evolutionary biology (Nei et al. 2010). One evolutionary context that has drawn increasing attention from evolutionary biologists is the context of ‘range expansions’. Range expansions are a ubiquitous phenomenon in nature involved in processes such as biological invasions (Parmesan and Yohe 2003; Walther et al. 2009), adaptive radiations (Rundell and Price 2009), speciations (Thorpe 1984; Hewitt 1996), pest and disease outbreaks (Jepsen et al. 2008; Roth et al. 2010), and post-glacial recolonizations (Hewitt 1996). Contractions and recolonizations following glacial oscillations are immensely common in nature, not only in temperate areas, but in tropical and subtropical regions, as well (Colinvaux et al. 2000; Hewitt 2000). Therefore, it is probably safe to say that range expansions are likely involved in the evolutionary history of most of the organisms on the planet.

In the *selection vs. neutrality* discussion, range expansions are particularly important because populations increasing their range tend to do so over environmental gradients, leaving room for selection to act, possibly leading to local adaptation (Hewitt 1996). When different forms are established across this gradient, a *cline* is produced (Endler 1977). Clines have been thoroughly studied in the context of hybrid zones, where two allopatric populations get into secondary contact forming a tension zone in which the hybrids are selected against, so that the width of the cline is inversely proportional to the strength of selection (Barton and Hewitt 1985). The same rationale was later applied to clines appearing in ecological transition zones (i.e. ecotones): Mullen and Hoekstra (2008), in what has become a classical example, demonstrated that strong selection maintains two color-morphs of deer mice separated in two different habitats. These studies, however, have concentrated on small geographical scale clines. When it comes to large-scale clines (such as those appearing across continents) the literature is relatively scarcer with some theoretical studies focused on gene frequencies (Bazykin 1969; Endler 1977) and quantitative traits (Barton 1999; Leimar et al. 2008), and other empirical studies mostly dedicated to the description of clinal patterns in organisms like *Drosophila* spp. (Hallas et al. 2002; Weeks et al. 2002), *Populus tremula* (Ingvarsson et al. 2006), *Quercus petrea* (Zanetto and Kremer 1995), *Pinus sylvestris* (García-Gil et al. 2003), *Arabidopsis thaliana* (Kronholm et al. 2012), and yet other plant species (Savolainen et al. 2007). However, no attempt to measure selection in any of these or any other large-scale systems has been carried out, to our knowledge.

Still in the context of expanding populations, Edmonds *et al.* (2004) proposed that the formation of (genotypic) allele-frequency clines across environmental gradients could also (and mainly) be caused by a purely neutral process during range expansion: the allele-surfing phenomenon, further studied and named by Klopfstein *et al.* (2006). In populations undergoing a range expansion, mutations arising at the front of the wave of expansion can “surf” on this wave and increase in frequency simply due to a series of founder events. This surfing leaves behind a pronounced cline in allele frequencies, which may in turn have an effect on phenotype, generating a phenotypic cline. Recent studies are bringing a growing body of evidence that allele surfing alone is capable of producing many of the allele-frequency clines observed in natural populations (Currat et al. 2006; Excoffier and Ray 2008; Hofer et al. 2009). Some more recent findings even show that range expansions might allow for the accumulation of deleterious mutations generating an ‘expansion load’ in populations of recently colonized areas (Peischl et al. 2013; Peischl and Excoffier 2015; Gilbert et al. 2017).

Expansion load has been demonstrated to have a complex interaction with the adaptation dynamics of an expanding population. The existence of an environmental gradient can reduce the accumulated genetic load during the expansion process, while the consequent maladaptation in the front-end of expansion may in fact reduce the speed of the process (Gilbert et al. 2017). Moreover, the steepness and patchiness of the environmental gradient combined with different genetic architectures can have significant consequences on the outcome of the range expansion, as well. In fact, if the gradient is too steep and the genetic architecture relying on large-effect alleles, the range expansion might fail altogether (Gilbert and Whitlock 2017).

The focus of this work is not on expansion load, and rather on the adaptive processes possibly involved in range expansions. And there is indeed evidence that range expansions may foster adaptive processes, bringing about the idea of adaptive clines. For example, White *et al.* (2013) found indications of adaptive evolution in an ongoing range expansion in Irish bank voles, where several genes related to immune and behavioral systems were shown to form consistent clines across three independent transects of the expansion. Empirical evolutionary studies have also suggested that range expansion could facilitate adaptive change (Gralka et al. 2016). Furthermore, it appears that dispersal ability itself is a trait commonly affected by selection in range expansions: higher dispersal is often selected for in the margins of an expansion, as theoretical analyses suggest (Travis and Dytham 2002). Empirical support for this finding has been documented in several species (Hughes et al. 2007; Monty and Mahy 2010; Moreau et al. 2011). Furthermore, rapid adaptation to climate variation also facilitates range expansion, as has been verified in the invasive plant *Lythrum salicaria* in North America (Colautti and Barrett 2013). The body of evidence favoring selection in range-expansion systems is substantial, and it often includes the examples of the (continental) large-scale clines mentioned above, as well (Bazykin 1969; Endler 1973; Barton 1999; Leimar et al. 2008). One particularly interesting case in large-scale cline and range expansion systems is the European barn owl (*Tyto alba*) and its coat-color cline (Antoniazza et al. 2010, 2014). In this species, a gradient of colors has established across Europe, probably during or after a post-glacial range expansion, with white morphs nearly fixed in the southwest and dark-brown morphs in the northeast. This and the above-mentioned cases all suggest selection has been acting. However, the current challenge persists in (i) distinguishing neutrality from selection and (ii) properly measuring the strength of natural selection in large-scale clinal systems involved in range expansions.

The question of whether or not one is able to assess selection in range expansions is still unsolved. Here, we take advantage of spatially explicit simulations to investigate the role of selection in the context of range expansions. First, we assess the ability of selection to leave a distinctive signature of its activity on the populations, despite the occurrence of the complicating demographic effects of range expansions (e.g. allele surfing). Second, with approximate Bayesian computation (ABC) (Beaumont et al. 2002), we address the detection and estimation of natural selection operating in this system. Finally, focused on the estimation of selection, we also explore the effect of other demographic and genetic parameters (nuisance parameters) on the accuracy of the selection estimates. Variations in these parameters may affect the probability of allele surfing. Therefore assessing the robustness of selection estimates across these parameters can bring valuable insight on the interplay between neutrality and natural selection in the ubiquitous demographic scenario of range expansions.

## Material & Methods

### Range expansion

Simulations were run in a rectangular world 5 patches wide and 51 (0-50) patches long (Fig. 1A) in an internal-development version of the program quantiNEMO2 (Neuenschwander et al. 2008a, 2018). To mimic a range expansion, only the left-most patches started the simulations occupied at their carrying capacity (K = 100). These five patches evolved without any range expansion for arbitrary 100 generations in order to establish a background of genetic diversity, mimicking a refugium. The colonization of the remaining patches occurred after this initial phase and lasted 400 generations, at a speed that depended on the migration rate (m, uniform [0.1, 0.4]) and the intrinsic growth rate of each patch (r, uniform [0.2, 0.8]). We further varied narrow-sense heritability value (h^2^, details below) and mutation rate (μ, log-uniform [10^−5^, 10^−2^]), which were used as “nuisance” parameters to test the robustness of the selection-related parameter’s estimates. As neutral genetic markers, ten multi-allelic loci were simulated with the same mutation rate implemented for the quantitative loci (μ) and a single-step mutation model, mimicking microsatellite markers.

**Figure 1:**
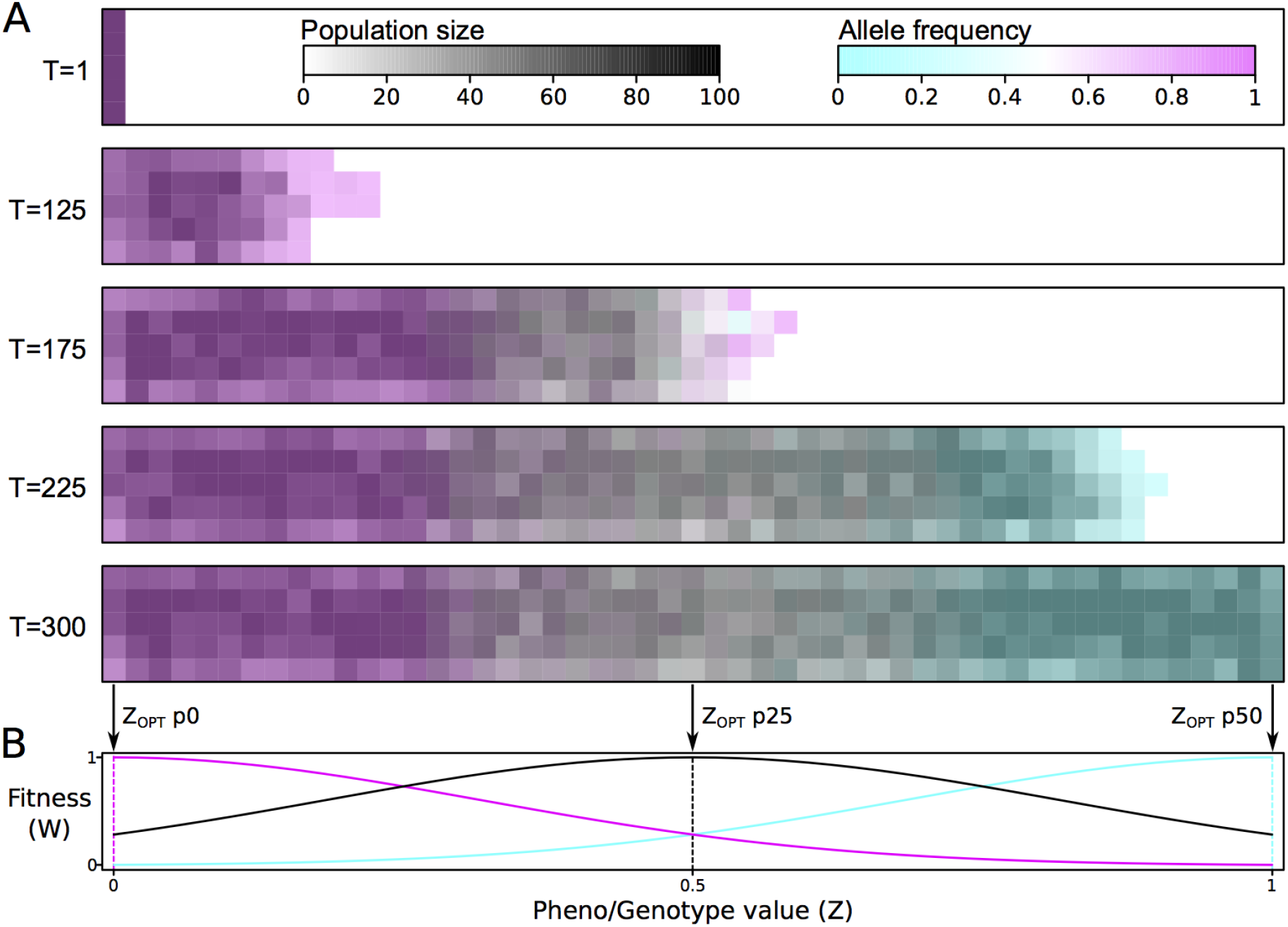
Implementation of the simulations with range expansion over a selection gradient. In **A**, the range expansion process over 300 generations (T), across the simulated map (51×5 demes). Two layers overlap here: population size (gray scale, underneath) and frequency of the allele adapted to the left-hand side of the map (cyan-magenta scale). In **B**, the fitness landscape for three patches from above (p0 magenta, p25 black, and p50 cyan) with selection intensity ω=0.1 and pheno/genotype space defined between 0 and 1. Note that the x-axis in B (Z-value) is different from the one in A (deme position p).

### Selection implementation

Fig. 1B illustrates how selection was implemented: we assumed a local hard stabilizing-selection scheme with a gradient of optima along the colonization path. On the left-hand side of the map, the selective optimum was defined at one extreme of the phenotypic range (Z_OPT_ = 0); while, at the right-hand side, it was set to the other extreme (Z_OPT_ = 1). Each patch along the colonization path had a different optimum value (Z_OPT_), linearly distributed between 0 and 1. Individual fitness is given by the function:

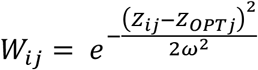

where W_ij_ is the fitness of individual i from patch j with phenotype Z_ij_, where the patch optimum is Z_OPTj_ and the selection intensity (identical for all patches) is given by ω. This latter parameter determines the strength of selection in our model (ω, log-uniform [0.1, 100], Fig. 2A). The ω parameter translates directly into a selection coefficient (s) (Fig. 2B) according to equation:

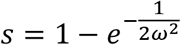

where s is the selection coefficient – defined as the difference in fitness between the two extreme pheno/genotypes (Z = 0 or 1) at any of the ends of the map – and ω is selection intensity, as already defined above. Part of the phenotype is environmentally determined, depending on trait heritability (h^2^). We explored a wide range of heritability values (h^2^, uniform [0.01, 1]), kept constant over time within the same simulation. Our goal is to estimate the selection coefficient (s), having nuisance parameters corresponding to the heritability of the trait (h^2^), migration (m), mutation (μ) and growth (r) rates.

**Figure 2:**
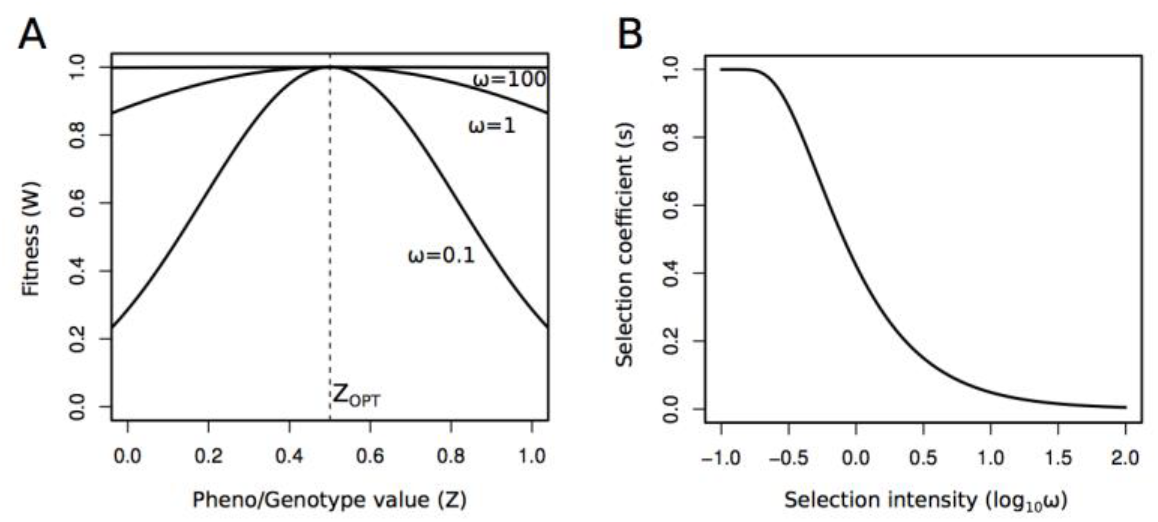
Fitness distribution and selection coefficient under different selection intensities (ω). In **A**, different fitness distributions with Z_OPT_ always at 0.5, as in patch p25 (see Fig. 1B), depicting the extremes of the ω prior distribution ω=0.1 and 100. In **B**, the effect of ω on the difference of fitness [i.e. selection coefficient (s)] between opposing pheno/genotype values at the extreme patches (p0 and p50).

### Six genetic architectures

Six different genetic architectures were implemented for the trait under selection where the allelic effects were entirely additive within and between loci. First, we assumed a trait encoded by one locus and two co-dominant alleles (1L2A). In this case, only one mutation was needed to make the leap between the two extremes of phenotype. The second model still involved only one locus, but with multiple alleles (1L10A), whose effects on the phenotype were linear and additive. Here, there are only two alleles completely adapted to the two extremes of the environmental gradient; all other alleles have intermediate values, which are able to match the intermediate optima along the colonization range. The third genetic architecture was that of a trait encoded by ten bi-allelic loci (10L2A) where all loci are required to adapt to obtain the extreme phenotypes. A second version of this architecture was one with the same number of loci and alleles, but with allelic effects large enough for a mutation at a single locus to allow for perfect adaptation to the extremes (10L2A+). A fifth architecture involved 10 alleles at 10 loci (10L10A), similar to 1L10A, but extended to ten independent loci. Similar to the extension of large allele effects applied in 10L2A+, a sixth architecture was defined with the possibility of any given locus as being able to modify the phenotype across its complete range (10L10A+). Mutation rate was scaled to the number of loci encoding the trait, so that the trait’s mutation rate was the same across architectures (i.e. it was 10× lower for each locus in the 10L architectures).

### ABC for selection

One suitable way to address complex evolutionary questions is to implement approximate Bayesian computation (ABC). With this approach, one can assess the probability of different scenarios and parameter values therein via summary statistics, thus dismissing the need of an exact likelihood function (Beaumont et al. 2002). Summary-statistic values are taken from the observation (i.e. the real populations) and compared to the values of the same statistics obtained in simulations. A large number of simulations are then used to explore different combinations of parameter values; the simulations that better match the summary statistics values of the observation are then used to draw a posterior distribution of parameter values. As a Bayesian method, ABC can (and should) incorporate prior information on the parameter distributions into the simulated model. Here, we applied ABC to the estimation of selection in a spatially explicit setting involving range expansions. Since this a simulation study, the observations were also taken from the simulations in the form of pseudo-observations (see below).

### ABC: summary statistics

Based on our previous experience with a similar set-up in natural populations of barn owls (Antoniazza et al. 2010, 2014), we decided to focus on isolation-by-distance (IBD) pattern statistics as the statistics more likely to reveal the presence of selection: From the correlation between pairwise geographic distance and pairwise pheno/genotypic distance, we extracted the mean, slope and sum of residuals for ten neutral multi-allelic markers (F_ST_), and the phenotype (Q_ST_). Finally, we also retained the difference of slopes of IBD between the phenotype and the neutral markers (Δ-slope), which represents how much steeper is the differentiation in the quantitative trait when compared to the one produced by the neutral loci (Fig. S1).

### ABC: parameter estimates and estimability assessment

We tested the precision and accuracy of parameter estimates through ABC’s validation approach as implemented in ABCtoolbox (Wegmann et al. 2010). Since the actual parameter values for all simulations are known (pseudo-observations), the ABC parameter-estimation pipeline was used to assess the quality of the estimates (i.e. how close the estimates were to the actual values). This was done by comparing 1000 of these estimates with their actual pseudo-observed values taken directly from the simulations, for each one of the genetic-architecture models. This procedure involved retaining the 1000 (out of ~1 million) simulations with summary statistics values closest to the pseudo-observation’s, and then to use locally weighted linear regressions to obtain the posterior distributions for the parameter estimates (Wegmann et al. 2010). The overall estimability of selection coefficient for the different architectures was assessed using the coefficient of determination (R^2^) of the regression between the true value of the parameter (pseudo-observation) and the parameter point estimate (given by the mode of the posterior distribution) (Neuenschwander et al. 2008b). Two other statistics were also used to assess estimability: the root mean square error (RMSE), which depicts the prediction errors of our model by means of the mean absolute differences between pseudo-observations and estimates (Wegmann and Excoffier 2010); and proportion of the estimated posterior encompassing the pseudo-observed value for 50% and 95% of the higher-posterior density intervals (proportion of HPD50% and 95%). This latter statistics may indicate a low accuracy, when proportion of HPD50% << 0.5, or HPD95% << 0.95; or excessive conservativeness, when proportion of HPD50% >> 0.5, or HPD95% >> 0.95. Ideally, HPD50% and 95% should be exactly 0.5 and 0.95, respectively.

Moreover, to assess the effect of the nuisance parameters (m, r, μ, h^2^) on the estimability of selection coefficients, a second test was devised in which the parameter space of each one of the nuisance parameters was restricted to ten quantiles. The estimations of selection coefficient were obtained only in that restricted space. For example, heritability (h^2^) varied randomly from 0.01 to 1 across all simulations. To test whether estimates of selection were robust to a predetermined h^2^ value, we separated the simulations in ten different sets according to different quantile intervals of the h^2^ prior distribution – e.g. the first interval includes the simulation in which h^2^ ranges from 0.01 to ~0.1. For each of the h^2^ intervals, we obtain estimations of selection coefficient (s) that were then compared to their pseudo-observed value. This was also done for the other three nuisance parameters (m, r and μ) and across all six genetic architectures. Quantiles of the parameter values, instead of fixed bins, had to be used in order to insure that all estimates were made based on the same number of simulations. This is because the inherent sampling process, combined with the failure of some simulations (see supplement), does not necessarily leads to the same density of simulations across the whole parameter space. So, for each quantile interval, 1000 estimates were run with 500 retained simulations, and the estimability was again measured by means of R^2^, allowing for comparisons across the quantiles.

## Results

Overall, the statistics related to the regressions between pairwise differentiation (Q_ST_) and pairwise geographical distances were very sensitive to variation in selection strength, regardless of the genetic architecture implemented (Fig. 3). In particular, the difference of Q_ST_ and F_ST_ IBD slopes (Δ-slope) showed to be particularly responsive to small selection coefficients, while mean differentiation on the phenotype (mean Q_ST_) was more sensitive to moderate and high selection coefficients. Additionally, as expected for independent neutral loci, the statistics related to F_ST_ alone did not vary with the selection coefficients (results not shown). For nearly all architectures, mean Q_ST_ showed a constant quasi-linear increase with higher selection coefficients (Fig. 3A). The only two exceptions were the 1L2A and the 10L2A+ (with large-effect alleles) architecture models. In fact, these two architectures showed very concordant responses also in the other statistics, such as Δ-slope (Fig. 3B). In both cases, one can observe a lack of points for high selection coefficient values (s > 0.5). Indeed, these simulations failed to colonize the entire habitat (further examined below in ‘Discussion’). Moreover, Δ-slope, for all architectures, reaches an asymptote when s > 0.4. This is because, when selection is very strong, even closely neighboring demes are highly differentiated (high Q_ST_). This leads to high mean Q_ST_, but limits (or even reduces) the values obtained for the slope of differentiation across the environmental gradient (Fig 3B). Noteworthy are also the similarities between 1L10A and 10L10A+.

**Figure 3:**
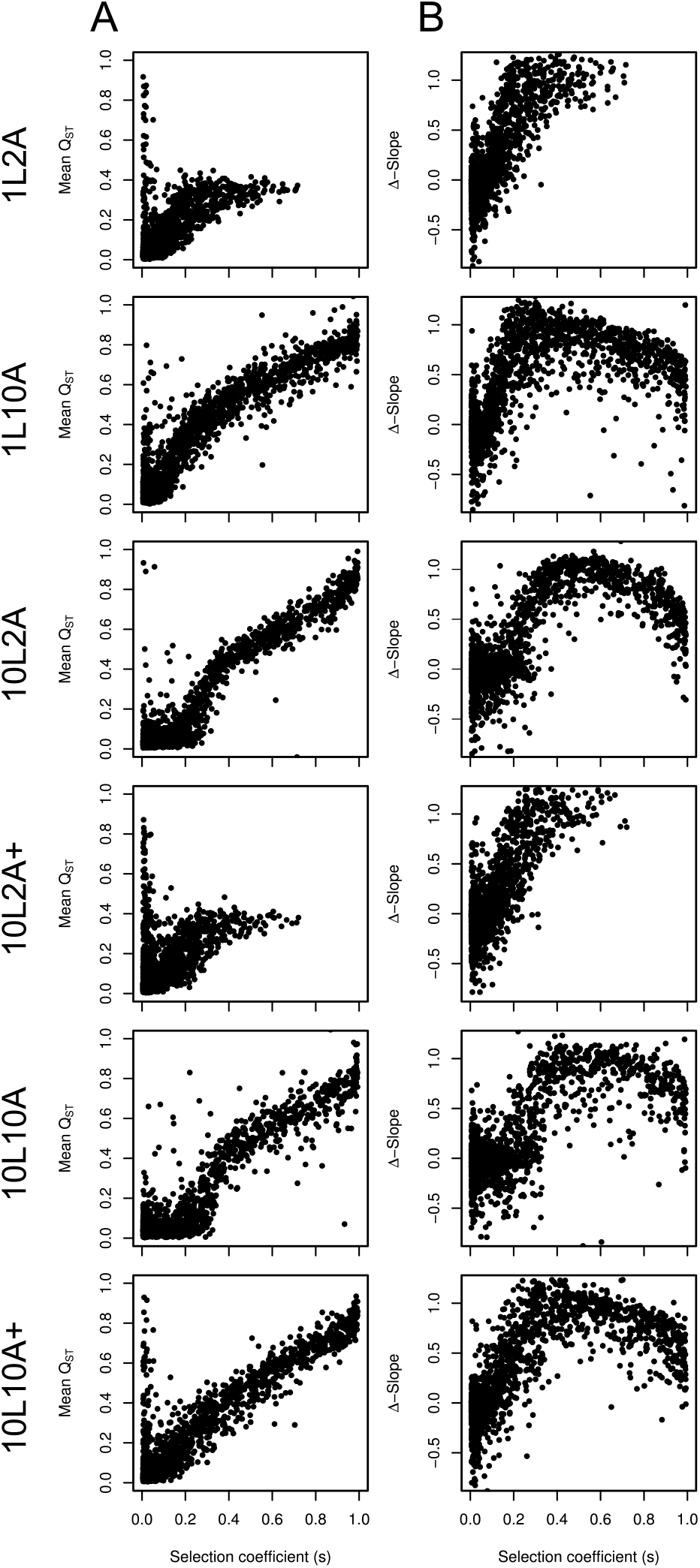
The relation between selection coefficient (s) and the most informative pattern statistics used to assess the selection coefficient. For all six architectures, in **A**, the response of mean differentiation across populations (Mean Q_ST_); and in **B**, the response of the difference between the Q_ST_ and the neutral F_ST_ slopes of IBD (Δ-Slope).

The quality of estimates for selection coefficient (s) in all models was high (Table 1, Fig. 4). The genetic-architecture models 1L10A, 10L2A, 10L10A and 10L10A+ had particularly high coefficients of determinations (R^2^ > 0.9), with 1L2A and 10L2A+ falling shortly behind (R^2^ > 0.7). This difference among the architectures derives from the differences in the summary statistics (above), where simulations with s > 0.5 failed to leave any signature on the summary statistics (Fig. 3), resulting in a limited range of s values (Fig. 4). Furthermore, the root mean square error values were proportionally low for all architectures (RMSE ≈ 5 to 9% of s estimates), implying a very high accuracy of estimates. The proportion of posterior-estimate distributions that encompassed the pseudo-observed value – both with HPD50% and 95% – resulted in conservative estimates (Table 1), with proportion values always larger than the HPD interval. This suggests that, even though accurate, the posterior distributions are not necessarily very precise, with rather wide ranges.

**Table 1:**
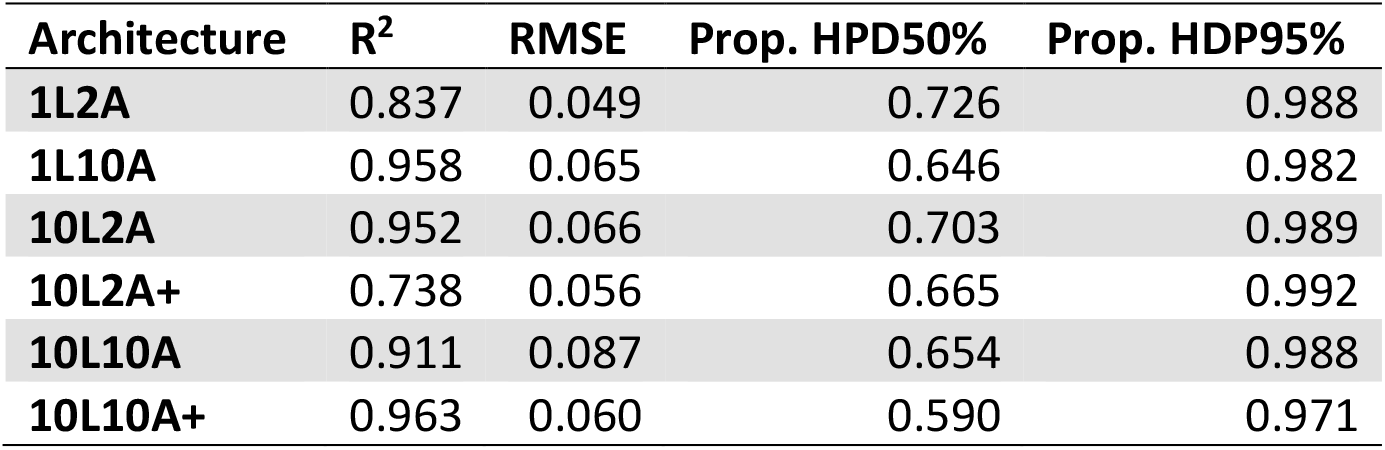
Assessment of selection coefficient (s) estimability for all genetic architectures. R^2^ stands for the coefficient of determination of the pseudo-observation on the estimates; RMSE is root mean square error of the estimates; and Prop. HPD50% and HPD95% represent the proportion of posterior distributions encompassing the pseudo-observed value. These values were obtained based on 1000 estimates, with 1000 retained simulations out of 1 million simulations, under a stabilizing hard selection system.

**Figure 4:**
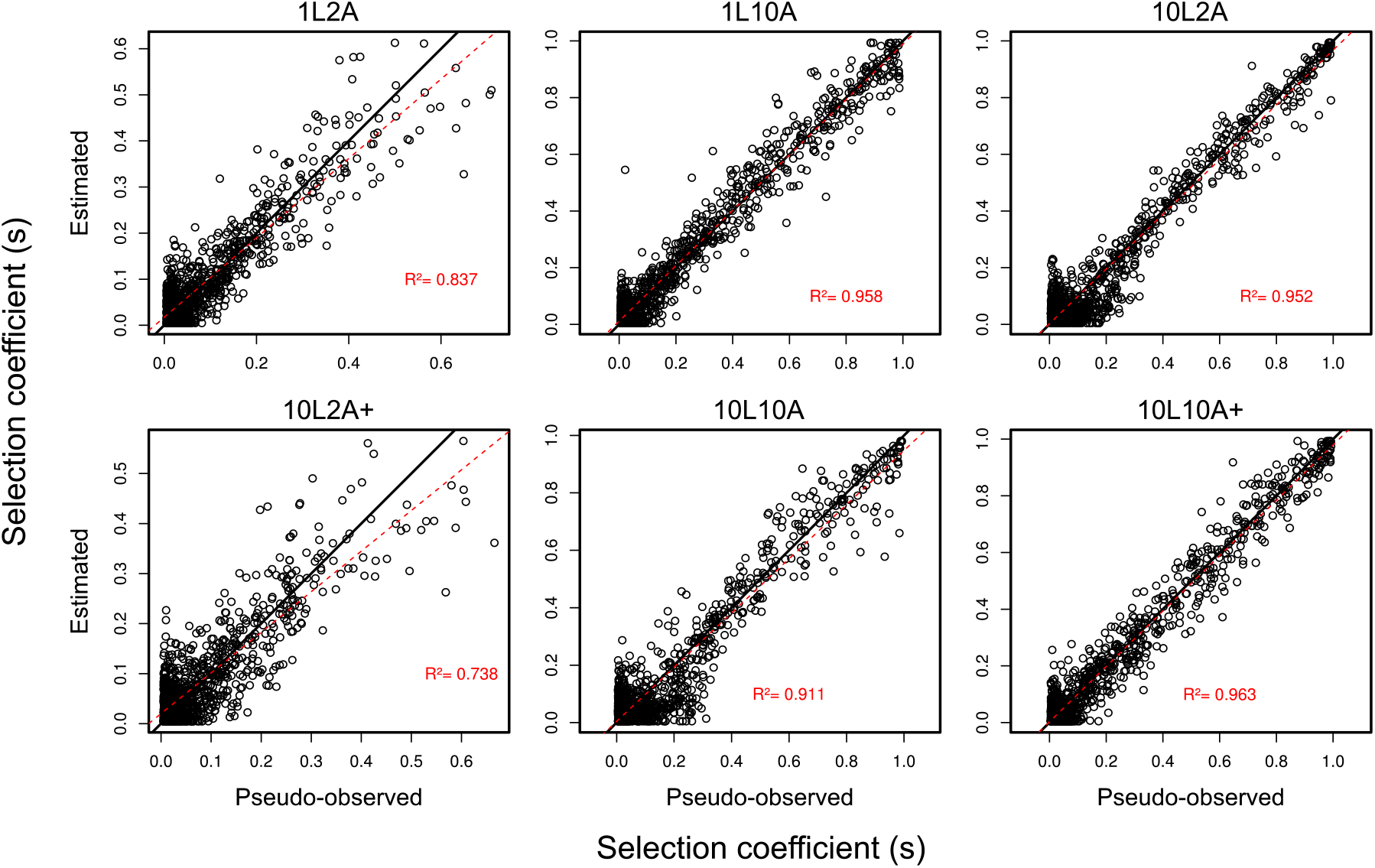
Validation plots, pseudo-observed vs. estimated, for selection coefficient (s). For each genetic-architecture model, a plot of 1000 simulations’ actual selection coefficient values (s) against their estimates (open circles). The back line stands for the perfect diagonal; and the red dashed line, the calculated linear regression. Coeffiecients of determination of the pseudo-observation on the estimates (R^2^) are also reported in red.

Remarkably, in our simulations, the estimability results are robust to the variation in the nuisance parameters and to the position in the largest part of the nuisance parameters’ space, with the clear exception of lower values of heritability (h^2^ < 0.1) for all architectures and also, to a lesser extent, lower values of mutation rate for some architecture models (Fig. 5). Interestingly, variation in migration (m) and growth rate (r) in the interval explored (m = [0.1, 0.4] and r = [0.2, 0.8]) has very little effect. Here too, there seems to be a ranking of estimation quality among the genetic-architecture models across the nuisance parameter quantiles: 1L2A and 10L2A+ being the worse (but still good); followed by 10L10A; and then having 10L10A+, 10L2A and 1L10A as the better ones.

**Figure 5:**
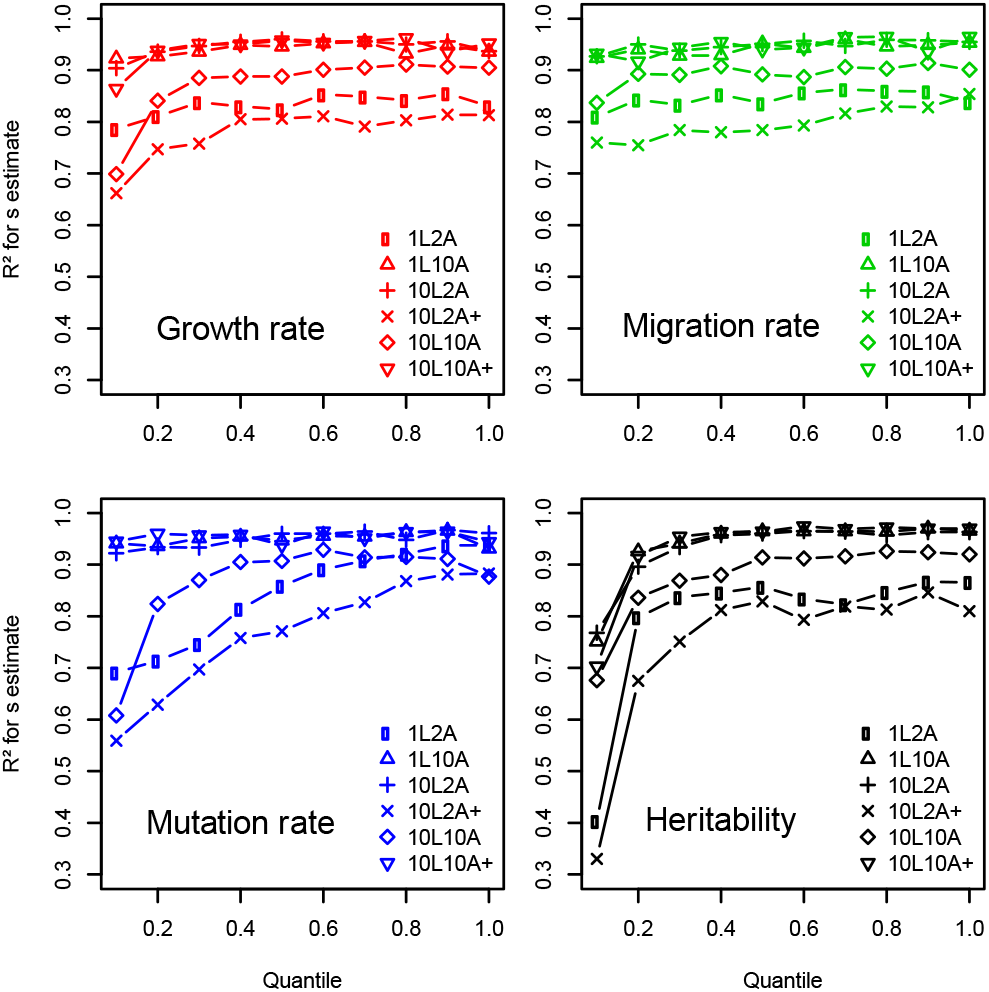
Estimability assessment across the nuisance-parameter space, for all genetic architectures. In each panel, the estimability of selection coefficient (by means of R^2^) is shown for ten different quantiles of the realized prior distributions fo the four nuisance parameters (each panel) and all six genetic architectures (within panels).

## Discussion

We have shown that it is possible to assess selection and estimate its intensity in range expansions by taking advantage of the information contained in IBD-derived statistics and by using spatially explicit simulations. Even though plenty of variance in the response of the summary statistics was observed when comparing the different genetic-architecture models, in all cases, selection left a distinctive signature on these statistics. It seems, however, that the probability of the populations to respond to selection was not the same across all architectures. In a nutshell, the more alleles and loci encoded the trait; the better was the estimation of the selection coefficient.

The architectures can be divided in three groups: (i) 1L10A and 10L10A+ with very high R^2^ and low RMSE (i.e. very good estimability), (ii) 10L2A and 10L10A with still high R^2^ and low RMSE values but with a distinct signature in the Δ-slope statistic (Fig. 3B), and (iii) 1L2A and 10L2A+ with slightly worse R^2^ and RMSE results. Not surprisingly, these last two architectures are also the ones that present the least number of allele combinations (within the phenotypic range between Z = 0 and

1) that could lead to adaptation across the selection gradient. The 1L2A model has only three possible genotypes to be translated into phenotypes. In essence, this architecture is just as capable as the others to adapt to the two extremes and the exact center of our simulated environment (patches p0, p50 and p25, respectively). However, this does not apply for any of the patches in between. In these other patches, there is no combination of alleles that would make an individual perfectly adapted to the local conditions. This same explanation applies to 10L2A+ because the large-effect alleles turn up to make too big a leap in between pheno/genotypic values (Z in Fig. 1). Indeed, if a second locus mutates as well in 10L2A+, the Z-value of the resulting phenotype would almost certainly fall outside the range of adapted phenotypes in all patches (Z = 0 to 1). This is why, when selection is too strong (s > 0.5), simulations failed to finish the colonization due to the recurrent extinction of pioneer populations. Conversely, all the other architectures present many more Z-value combinations allowing to locally adapt to all patches across the colonization range. These results may suggest that adaptation may be easier to occur when many loci and alleles contribute to a trait – offering several to many combinations of loci and alleles in order to adapt to the local conditions – in agreement with previous studies (Le Corre and Kremer 2012; Gilbert and Whitlock 2017).

It is important to highlight the impact of the inclusion of spatial information in the understanding of the effect of selection in range-expansion scenarios. The process of range expansion is essentially a spatial phenomenon and, to fully understand its outcome, a spatially explicit approach is warranted. Even though some of the statistics we used – mean F_ST_ and Q_ST_ – do not explicitly contain spatial information, it was only with the addition of Δ-slope and the other IBD-associated statistics that we managed to grasp the full extent of the of the effect of selection in range-expansion processes. The importance of the spatial dimension in population genetics is not a novel idea, though. It has been explored in numerous previous publications, both in the disciplines of phylogeography (Avise et al. 1987; Diniz-Filho et al. 2008) and in landscape genetics (Manel et al. 2003). Studies looking for signatures of selection, however, have been systematically neglecting the relevance of the spatial distribution of genes and phenotypes (Li et al. 2012).

Furthermore, combining more than one pattern statistics (at least mean Q_ST_ and Δ-slope, Fig. 3) seems to be of key importance to properly assess the effect of selection on populations facing range expansions. For instance, the analysis of mean Q_ST_ alone could lead to false positives when selection is very low (virtually zero), given that a few observations of high overall differentiation appear in these quasi-neutral conditions (Fig. 3A). Also, looking at Δ-slope alone could lead to false negatives – or simply lack of information – when selection is too strong, leading to less steep slopes than the ones observed at intermediate selection coefficients (Fig. 3B). Therefore, to properly benefit from our proposed ABC approach, we believe that one should always, of course, consider all available information contained in the different IBD pattern statistics.

Even though we modeled selection via intensity of selection (ω) – a parameter widely used in quantitative genetics (Falconer and MacKay 1996) – we decided to estimate selection through selection coefficient (s), which is a relatively more common measure in population genetics (Hartl and Clark 2007). Selection coefficient is a parameter whose effect on fitness (W) is directly accessible (W = 1 - s), making biological interpretation easier. Also, while ω had to be treated in the logarithmic scale (to obtain a more linear relation with the summary statistics), s could be dealt with in a linear scale. Besides, the results for estimability calculated for log_10_ω showed only a slight trend to lower R^2^ values and did not differ substantially from the ones obtained with s (Table S1). Regarding the scale of selection coefficient here, it is worth to remember that it concerns the difference in fitness in the extreme patches and the difference in fitness between the extreme pheno/genotypes (p0 and p50, Fig. 1). It becomes smaller as one approaches the center of the map and/or compares closer pheno/genotypes, and therefore represents the maximum strength of selection operating in the system.

We mentioned that some simulations “failed to finish the colonization altogether”. This requires further explanation. By failed simulations, we do not necessarily mean simulation where the population went extinct, but actually simulations that resulted in missing-data (NA) for any of the statistics. First, the simulations were run assuming a hard-selection system (i.e. individual fitness is absolute). So – even though local populations could react to loss of individuals via population growth – if selection was too strong and no locally-adapted individuals were yet present at the population, that specific deme would go extinct delaying or stopping the wave of expansion. Alternatively, we also ran the same simulations with a soft-selection system (supplementary material). These showed a lower failure rate, but did not affect further results, suggesting that the approach presented here is also robust to the softness of the selection implemented. Second, some architecture models lead to higher failure rates than others, predominantly due to the non-colonization effect described above. This is again related to the limited combinations of loci and alleles observed in architectures 1L2A and 10L2A+. As a result, the realized prior distribution (i.e. the parameter distribution after the removal of simulations containing NAs) for selection intensity (ω) – and therefore selection coefficient (s, as in Fig. 2) – was altered for these two architectures, being limited to ω = 10^−0.5^ to 10^2^ (s ≈ 0.8 to 0, respectively, Fig. S2 and S3). Beyond selection strength, for the other simulation parameters (i.e. nuisance parameters), there was no differential effect of the architecture model on the way these parameters produced simulations containing missing data. There was, however, a more elevated missing data production, for all architectures, associated with low mutation rates (when μ < 10^−4^), when not enough variation was produced to adapt to new environments; low growth rates (r < 0.3), when the negative effect of higher selection coefficients was stronger on the populations; and, to a lesser extent, higher migration rates, where the homogenizing effect of migration more often erased the differentiation signatures created by selection. As a result, the prior distributions for the nuisance parameters were altered after the removal of such failed simulations (Fig. S2). Consequently, the ten quantiles presented in Fig. 5 do not necessarily represent 10% intervals of the original prior distributions, but rather regular intervals taken from realized prior distributions. The analysis was done this way in order to have the same number of simulations out of which to make the estimates in each interval, allowing for a balanced comparison of estimability across quantiles.

The estimability of selection was little affected by variation in the nuisance parameters, as R^2^ remained well above 0.7 for all genetic architecture models across most of these parameters’ distributions. Some of the architectures seemed to be more sensitive to the noise caused by these parameters than others: Again, 1L2A and 10L2A+ showed to be the most sensitive models, probably, due to the lack of possible genotypic combinations, limiting adaptation to intermediary positions across the environmental gradient, as discussed above. However, the variation in mutation rate also had some effect on these architectures. The lower the mutation rate, the harder to deal with very strong selection, especially when combinations are limited. Another architecture in which selection estimability strongly responded to mutation rate was 10L10A. Curiously, this is the one with highest number of genotype combinations. This can be explained by the fact that it also is the architecture that needs the most mutations in order to adapt to the opposite environmental conditions during the range expansion. All ten loci need to adapt by fixing one of ten possible alleles each. Finally, as one could already expect, low values of heritability led to lower estimability for all architectures. Clearly, if the trait under selection has a very small genetic component, selection can do very little to affect the differentiation of the quantitative trait, leaving no signature of adaptation in the pattern statistics we explored, or any other statistics one could think of, as well.

It is still computationally expensive to run the individual-based spatially explicit simulations required to study the evolution of quantitative traits in range expansions, especially with several models of genetic architecture (e.g. ~350 CPU days for 1 million simulations on a Linux server with 2.4GHz Intel Xeon processors). This is because an ABC implementation generally requires many simulations (at least 1 million) to obtain reliable parameter estimates (Fagundes et al. 2007; Neuenschwander et al. 2008b), even though this can dependent at a large extent on the number of the parameters to be estimated (i.e. the dimensions of the parameter space to explore). Alternatively, improvements on the ABC algorithm such as MCMC-ABC (Wegmann et al. 2009) can help reducing the number of simulations needed for investigating a given question. Besides, selection was not the only parameter varying in our model. Nuisance parameters, even though not estimated, also affect the parameter space to be explored by the simulations. These do not have to be used, though: We added them to our analysis to assess the robustness of our estimates, but this does not need to be done in empirical studies. An approach that could be followed in such studies would be a two-step ABC analysis (Bazin et al. 2010), where (i) one would determine a neutral demographic background based on neutral markers and coalescent simulations, and (ii) then use the estimates of this previous step as priors for the following one in which individual-based simulations would be run to explore a different set of fewer parameters (e.g. selection coefficient and heritability), assuming that the effects of selection on demography would have already been captured in the first step.

Contrary to an impression one might get reading some of the recent theoretical literature on range expansions (Klopfstein et al. 2006; Travis et al. 2007; Excoffier et al. 2009; Peischl et al. 2013), selection is able to operate in such scenarios. Recent empirical studies have been showing evidence that adaptation has occurred in several cases (Hughes et al. 2007; Antoniazza et al. 2010, 2014; Monty and Mahy 2010; Road et al. 2012). When compared to allele surfing, selection seems to be much more efficient in producing differentiation across the range of an expansion, according to our results. Even though we observed consistent isolation by distance in the neutral loci (proxy for pure allele surfing), this isolation was always much lower than what was observed for the trait under selection.

The direct observation of some simulations provided evidence that locally maladapted variants could appear and reach relatively high frequencies during the range expansion process (Fig. S4), but these events tended to be transient and were quickly erased by selection, leaving virtually no signature after the whole map had been occupied. This observation may be the result of the model implemented here, where only one locus or a few loci were involved with selection and, therefore, could bear locally maladaptive (deleterious) variants. Another theoretical study, focused on the evolution of genetic load, provided evidence that, when many loci are involved, the overall deleterious load of populations undergoing range expansions tends to increase (Peischl *et al.* 2013). Indeed, there seems to be a decrease in the efficiency of purifying selection in purging a genome-wide deleterious load during range expansion (i.e. expansion load). However, here we investigated a process involving positive selection acting on one specific phenotypic trait whose genetic architecture was relatively simple. It is in this situation, we showed that natural selection during range expansions is still effective. Furthermore, in real populations, the simultaneous occurrence of adaptation at a given trait with the accumulation of an expansion load is perfectly possible and may be one explanation for the success of so many range expansions observed in nature. The combined effect of these two processes, however, remains to be more carefully investigated in the future.

Even though neutrality (including background selection) (Kimura 1984) should always be the null hypothesis for any investigation of a process leading to a given observed pattern, we believe that here we have gathered sufficient *in silico* evidence that selection can operate on range expansion scenarios, leaving a distinguishable signature in spatially explicit statistics. Furthermore, this signature allows estimating the strength of selection operating on the study system and could be promptly used in empirical studies investigating selection in range expansion scenarios – which could be post-glacial recolonizations, species invading new habitats, or populations coping with environmental changes. All of these processes were and still are very common, not only in temperate regions (Hewitt 2004), but also anywhere else on the globe, rendering the spatially explicit ABC approach presented here particularly valuable.

## Acknowledgments

The computations were performed at the Vital-IT (http://www.vital-it.ch) Center for high-performance computing of the SIB Swiss Institute of Bioinformatics. We would like to thank Laurélène Faye and Rémi Matthey-Doret for helping in initial stages of this project and Sylvain Antoniazza for many fruitful discussions that helped in the development of this study. JG had financial support from the Swiss National Science Foundation (SNSF) grant number 31003A_138180.

